# The prevalence of two known pathogens of Atlantic Blue Crab *Callinectes sapidus* (Rathbun, 1896) in coastal Adriatic, Aegean sea, and Atlantic Iberian coast

**DOI:** 10.1101/2025.02.14.637520

**Authors:** Eric Schott, Marina Brailo Šćepanović, Jasna Maršić-Lučić, Olivia Pares, Mingli Zhao, Jerko Hrabar, Luka Glamuzina, Marijana Pećarević, Sanja Grđan, João Pedro da Silva Encarnação, Pedro Morais, Ana Pešić, Ilija Ćetković, Branko Glamuzina

## Abstract

In the Mediterranean Sea, the abundance of the invasive portunid crab, *Callinectes sapidus*, has dramatically increased in recent years. This raises concerns about damage to ecosystems, but also offers opportunities for exploitation of a new fishery. Newly invasive species may escape from pathogens in their native range, may introduce new pathogens, or can become host to endemic pathogens. Understanding these factors is important for predicting or managing natural resources in the invaded range. This study investigated the prevalence of two pathogens common in *C. sapidus* in its home range of North America: the reovirus CsRV1 and the protozoan parasite *Hematodinium perezi*. In crabs collected from Aegean, Adriatic, and Atlantic waters, the CsRV1 virus was not detected. In contrast, the parasite *H. perezi* was found in crabs from all areas except the Aegean Sea. Sequence analysis of the *H. perezi* ITS1 gene indicated that the strains observed are most related to genotypes already described in Europe and the Mediterranean, and not to strains from the Americas or Asia. The arrival of new species and new potential pathogens is ongoing through transfer of ballast water to the Atlantic and Mediterranean. Although systems are in place to exchange or inactivate ballast water, it is advisable to continue and expand surveillance for pathogens in introduced species, to inform management of movement of these species between regions.

## Introduction

The Atlantic blue crab *Callinectes sapidus* Rathbun,1896 (Brachyura: Portunidae) is a swimming crab native to the western Atlantic in coastal and estuarine waters of New England (USA) to Uruguay, including the Gulf of Mexico and the Caribbean Sea (Mancinelli *et al*., 2021, 2017). In recent years it has also emerged as one of the most concerning invasive species in the Mediterranean Sea. It was first recorded (as *Neptunus pelagicus*) in Europe in 1900 (Bouvier 1901), and appears to be established on the Atlantic coast of France and England (Pezy *et al*., 2019). Other records show the species has also moved south to the upper Iberian coast (Ribeiro and Veríssimo 2014). In the Mediterranean Sea it was first recorded in 1949 in the Gulf of Venice, Italy, in the northern Adriatic Sea (Giordani Soika 1951), and possibly as early as 1935 near Turkey (Enzenroß *et al*., 1997). It has become invasive in the eastern-most and western basins of the Mediterranean and the Black Sea (Vella *et al*., 2023). Recently, the species was reported in Malta and Sicily (Vella *et al*., 2023), Sardinia (Culurgioni *et al*., 2020), Spain (Clavero *et al*., 2022), the Adriatic and Aegean seas (Glamuzina *et al*., 2023, 2021; Cilenti *et al*., 2015; Kevrekidis *et al*., 2023), and the Mediterranean coast of Africa (Kara and Chaoui, 2021; Taybi and Mabrouki, 2020). Also recently, *C. sapidus* has appeared in abundance on the southern Atlantic Coast of Portugal, and has been reported on the Atlantic coast of Morocco, possibly reflecting an expansion through the straits of Gibraltar (Morais *et al*., 2019; González-Ortegón *et al*., 2022; Lamkhalkhal *et al*., 2024).

The life history of *C. sapidus* includes a 28-35 day oceanic planktonic phase, before metamorphosis to benthic postlarvae (megalopa stage), that contributes to dispersal. Blue crabs are relatively short-lived, with an estimated longevity of 2-3 years in some parts of the US native range (Kahn and Hestler 2005), and a recent estimate of 4.8 years in an Italian lagoon (Mancinelli *et al*., 2024). Therefore, in locations with consistently high crab abundance, there must be recurring sources of larvae, either through local self recruitment or from broodstock sources upstream in oceanic currents. For example, in the south-eastern Adriatic Sea, a strong anti-clockwise gyre connects the coast of Italy and Croatia, while in the Aegean Sea, where blue crabs have been abundant for decades (Holthuis 1961) currents are weak and the many islands ensure promote connectivity and self recruitment (Estournel *et al*., 2021). The local-recruitment hypothesis is supported by Schubart *et al*., (2022) who found low genetic connectivity between different populations of *C. sapidus* in the invaded range, consistent with multiple founder introductions.

The success of the blue crab in new regions is likely enhanced by its omnivory and ability to adapt to a wide range of salinity, temperature and dissolved oxygen (Glandon *et al*., 2019). Success of non-native species in a new range is often also attributed to an escape from predators and infectious diseases that suppressed their numbers in the home range (sometimes referred to as the enemy release hypothesis) (Blakeslee *et al*., 2009; Torchin *et al*., 2007; Schultz *et al*., 2019). While a lot has been reported on occurrence and the invasiveness of *C. sapidus* in the Mediterranean, there is still little knowledge of the infections that it may acquire in this new range. The enemy release (pathogen escape) concept assumes that newly arrived species have not carried their home-range pathogens with them.

In its North American range, *C. sapidus* is infected by a number of pathogenic parasites and viruses (Shields and Overstreet 2007). The most studied is the parasitic dinoflagellate, *Hematodinium perezi*, which is known to cause mortality in *C. sapidus*, and is a host generalist that infects a wide range of crustacean species. The type species is described in *Liocarcinus depurator* from the Atlantic coast of Europe (Chatton and Poisson, 1931; Small et al. 2102).

Three different molecular *H. perezi* clades are recognized, each characteristic of different global regions, and distinguished by genomic changes in the ribosomal RNA gene ITS1 region (Small *et al*., 2012, Pagenkopp-Lohan *et al*., 2012). The genotype infecting crabs in the Mediterranean and Atlantic coast of Europe is referred at Type I, while those in Asia, often associated with mortality in aquaculture, are type II, whereas *H. perezi* found in *C. sapidus* and other hosts on the Atlantic estuaries of north America are type III (Small *et al*., 2012).

In the United States, regional population declines of *C. sapidus* have coincided with elevated prevalence *Hematodinium* sp. (Lee and Frischer 2004). A related *Hematodinium* sp. that infects Norway lobster *(Nephrops norvegicus*) is a known driver of mortality in this fishery on the west coast of Scotland (Stentiford and Neil 2011; Molto-Martin *et al*., 2024). In the Mediterranean, there are few reports of *H. perezi* (or other *Hematodinium* sp.) in any host, however it has been reported in *C. sapidus* and other crabs in the south of Italy (Pagliara and Mancinelli 2018; Mancinelli *et al*., 2013).

The life cycle of *H. perezi* is complex, with a progression of several morphological cell types within the host, culminating in the release of dinospores that emerge from mouth or gills (Stentiford and Shields, 2005). Transmission of the parasite is thought to be via dinospores, which remain infective for hours to days in the water column (Coffey *et al*., 2012). Prevalence and infectivity of *H. perezi* are responsive to salinity, with higher prevalence in crabs captured in water above 18 psu, and dinospores becoming inactive at salinity below 10 psu (Messick and Shields 2000; Coffey *et al*., 2012).

The blue crab is also infected across its native range by a virus known as CsRV1, which was first observed in electron microscopy as “reo-like virus” (RLV) in the 1970s (Johnson 1977). This virus is associated with the majority of mortality in captive blue crabs in the United States (Bowers *et al*., 2010; Spitznagel *et al*., 2019). Surprisingly, the CsRV1 genome has a region of high genetic similarity to a reovirus, named P virus, described in the portunid crab *Liocarcinus depurator*, on the Mediterranean coast of France (Flowers *et al*., 2016; Montanie *et al*., 1993; Walton *et al*., 1999). There has been no publication on evidence of CsRV1 or P virus in the Mediterranean in the intervening 25 years. CsRV1 has been studied thoroughly within the crab’s native range (Zhao *et al*., 2020). CsRV1 is present from Uruguay to New England, but has not been reported in wild capture of any other species than *C. sapidus* (Zhao *et al*., 2020). The genotype of CsRV1 varies in a manner consistent with geographic origins (Zhao *et al*., 2023).

As *C. sapidus* continues to expand and increase in abundance across the Mediterranean and Atlantic coast of Europe, several authors point out that there are still considerable gaps in our knowledge of the ecology of this crustacean in the new range, even as the species is being exploited as a fishery resource in some locations (e.g., Mancinelli *et al*., 2017; Vella *et al*., 2023; Azzurro *et al*., 2024; Galil 2024). Given the importance of *H. perezi* and CsRV1 in the ecology of *C. sapidus* in its native range, it is relevant to investigate how these pathogens are distributed in *C. sapidus* in the invaded range. Given the global distribution of *H. perezi* and other *Hematodinium* sp., the bi-hemispheric distribution of CsRV1 and molecular evidence that CsRV1 may have a close relative in Europe, we investigated the prevalence of these two known *C. sapidus* pathogens in the Adriatic and Aegean Seas, where *C. sapidus* has been reported for over a decade, and on the newly-invaded coast of Portugal in the Guadiana estuary.

## Materials and methods

### Crab sampling and environmental data

Thirty adult *Callinectes sapidus* were collected from each of 6 estuaries in the Mediterranean and Atlantic Iberian coast between 2021 and 2023 (Figure 1), for a total of 180 crabs.

**Figure 1.**
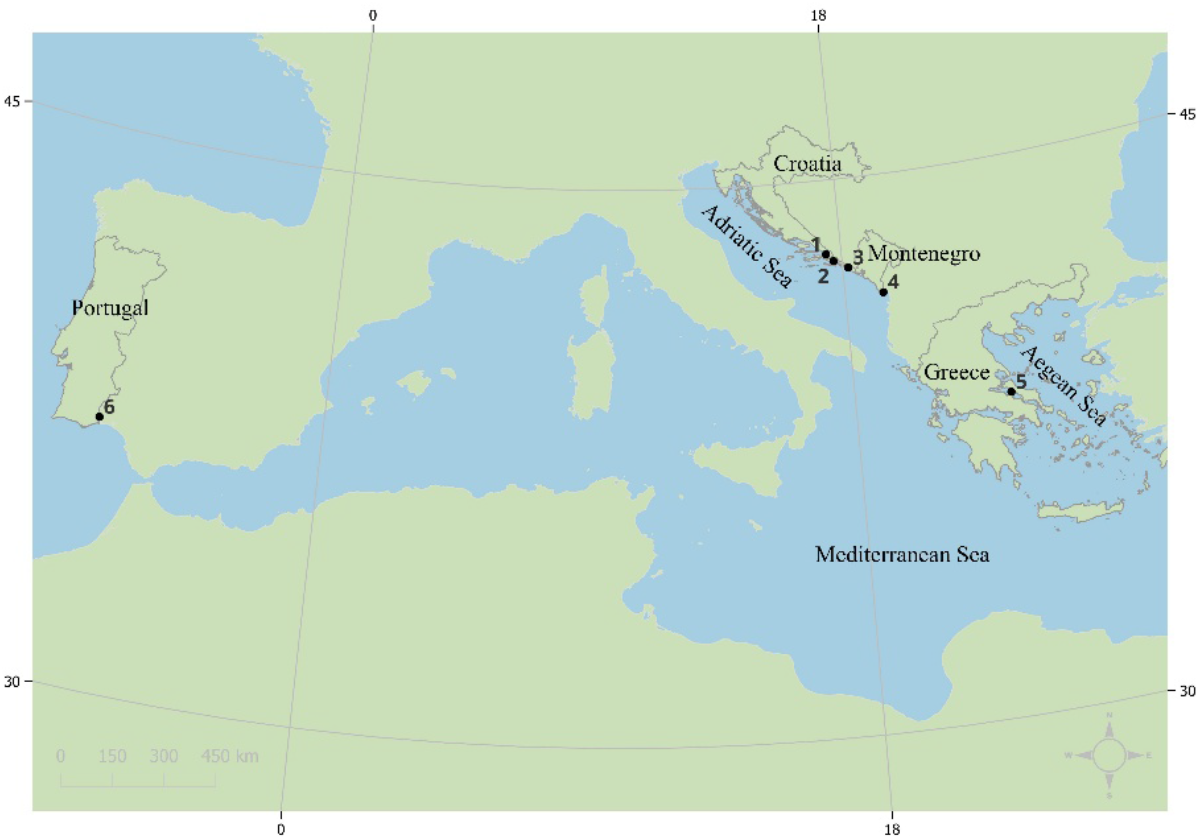
Sampling locations on the Adriatic, Aegean and Iberian Atlantic coasts. Numbers at each location correspond to numbers in Table 1.

Wire traps were used in Mediterranean sampling locations of the South Adriatic Sea in Croatia (Parila lagoon in Neretva river delta, Ston saltworks, Mlini in Župa Bay) and Montenegro (Ada Bojana). Blue crabs originating from the Aegean sea (North Euboean Gulf) in Greece were also captured using commercial wire traps and obtained from the fish market. Mediterranean crab samples were transported to laboratory on ice and processed immediately upon arriving.

**Table 1.** *Callinectes sapidus* sampling sites, dates and environmental conditions.

| Location | Map ID | Collect date<br>(month-year) | Latitude<br>(°N) | Longitude<br>(°E) | Temp<br>(°C) | Sal<br>(psu) |
| --- | --- | --- | --- | --- | --- | --- |
| Parila lagoon,<br>South Adriatic Sea | 1 | February and March<br>2021 | 43.0307 | 17.4512 | 15 | 18 |
| Mlini,<br>South Adriatic Sea | 2 | February and May<br>2021 | 42.8301 | 17.6975 | 21.7 | 38 |
| Ston saltworks,<br>South Adriatic Sea | 3 | May, June and<br>October 2021 | 42.6178 | 18.2031 | 18.8 | 38 |
| Ada Bojana,<br>South Adriatic Sea | 4 | September 2022 | 41.8538 | 19.3572 | 22.5 | 32 |
| North Euboean Gulf,<br>Aegean Sea | 5 | April 2023 | 38.7107 | 23.2851 | 15.0 | n.a.* |
| Foz de Odeleite,<br>Guadiana estuary | 6 | June 2023 | 37.3534 | -7.4409 | 18.3 | 13 |

\*n.a. not available

Temperature and salinity were measured at the time of the sampling in Parila lagoon and Ston saltworks by YSI Pro30 probe, and in Ada Bojana by HANNA HI 98194 multiparameter probe. Environmental data for Mlini were acquired from a year round study of the Župa bay (M. Pećarević, personal communication) while the temperature of the Aegean sea was obtained by daily satellite readings provided by NOAA, some 70 km southeast from the sampling site.

Crabs from Atlantic coast were captured by fishermen using gill and fyke nets in the tidal Guadiana River in Portugal (bordering Spain). Salinity and temperature were collected as a contribution to the International Group for Marine Ecology Time Series (IGMETS) monitoring program, in the Guadiana lower estuary station, southwest Iberian Peninsula. These crabs were transported to shore within 2 hours, and frozen at −20 °C until processing in the laboratory.

### Dissection, nucleic acid extraction and polymerase chain reactions

Crab legs (fresh or partially thawed on ice) were externally cleaned with RNaseZap (Thermo Fisher Scientific) and dissected with new razor blades. Two samples of 50-100 mg muscle tissue were dissected from a walking leg of each crab and stored at 4°C in 96% ethanol (Gram-Mol, Croatia) for DNA analysis, as well as in TriReagent solution (Thermo Fisher Scientific) for RNA analysis. All samples were transported to the Laboratory for Aquaculture of the Institute of Oceanography and Fisheries in Split, Croatia for nucleic acid extraction and histological analysis see Methods 2.3.

#### DNA extraction

Genomic DNA was extracted following a simplified phenol-chloroform-isoamyl DNA isolation procedure (Laird *et al*., 1991). Resulting DNA pellets were suspended in 30 µl of TE buffer, checked for purity and concentration by NanoPhotometer (Implen) and stored at −20°C until shipment. DNA was then re-precipitated with ethanol and shipped as precipitates to the Institute Marine and Environmental Technology (IMET) in Baltimore, USA for PCR analyses.

#### RNA extraction

For the RNA extraction the tissue was homogenized with 1 mL TriReagent solution (Thermo Fisher Scientific) with ceramic beads using MagNa Lyser (Roche) and then the procedure followed the protocol used by Spitznagel *et al*. (2019). Obtained RNA pellets were suspended in 50 µl of nuclease-free water and checked for purity and concentration by NanoPhotometer (Implen). RNA was re-precipitated in ethanol and pelleted nucleic acid samples were shipped to the Institute of Marine and Environmental Technology (IMET) in Baltimore, USA where they were stored at −20°C until resuspension in nuclease-free water for RT-qPCR analysis.

#### RTt-qPCR detection of CsRV1

The presence of CsRV1 RNA was investigated using quantitative reverse transcription PCR (RT-qPCR) methods adapted from Flowers *et al*., (2016), using a primer pair designed to amplify a 158-bp region of the ninth genome segment of CsRV1: 5’-TGCGTTGGATGCGAAGTGACAAAG-3’ (RLVset1F) and 5’-GCGCCATACCGAGCAAGTTCAAAT-3’ (RLVset1R). Positive controls and standard curves consisted of a 10-fold dilution series of purified double-stranded RNA (dsRNA) purified from crabs infected with CsRV1, and containing 10 to 10^6^ CsRV1 genome copies per μl, serially diluted in 25 ng μl^−1^ yeast tRNA carrier. The RT-qPCR cycling conditions and reagents were as described by Spitznagel *et al*. (2019), using qScript XLT One-Step RT-PCR Kit (Quantabio, Beverly MA) in 10 μl reactions containing 0.5 μM of each primer, on a QuantStudio 3 Real Time Thermocycler. Gene target copies were calculated as copies mg^−1^ of crab muscle, and samples with >500 copies mg^−1^ were recorded as CsRV1-positive according to Flowers *et al*. (2016), which is based on empirical measurement of background contamination from field and laboratory procedures. RNA amplification quality was assessed by using endpoint RT-PCR to amplify a fragment the blue crab small subunit ribosomal RNA transcript using primers CrabSSUfor 5’-CATGTCTAAGTACAAGCCGAA-3’ and CrabSSURev 5’-GGGTAATTTGCGTGCCT-3’, which produce a 405 nucleotide amplicon (GenBank JAHFWG010003967; Bachvaroff *et al*., 2021). PCR products were assessed by gel electrophoresis. RNA samples that failed to amplify with this rRNA primer set were deemed low quality and were not included in the CsRV1 analyses.

#### PCR detection of Hematodinium sp

The presence and amount of *Hematodinium* sp. DNA was assessed by the qPCR method of Nagle *et al*., (2009), which amplifies a 121-nucleotide region of the small subunit ribosomal RNA gene. Analyses were performed in duplicate on a QuantStudio 3 Real Time Thermocycler using PerfeCTa SYBR Green FastMix (Quantabio) with the following thermocycling conditions; 50°C for 2 min, 95°C for 10 min, followed by 40 cycles of 95°C for 15 s and 60°C for 1 min. The qPCR reactions included negative (no template) and positive (cloned target gene) controls. The number of *Hematodinium* sp. target copies was quantified by comparison to a standard curve (10 to 10^6^ copies per µl) of cloned *Hematodinium* sp. DNA (pES103, see Hanif *et al*., 2013) that was run on each plate. Based on the fraction of total DNA used in the qPCR reaction and the weight of the tissue from which it was extracted, the *Hematodinium* sp. infection intensity was expressed as the number of gene copies mg^−1^ of muscle.

#### Sequencing and analysis of H. perezi ITS1 gene

A 783 bp region of the 18S rRNA gene, from the SSU across the first internal transcribed spacer (ITS1) region to the 5.8S region, was amplified and sequenced for phylogenetic analysis. Amplification of this region was performed using the primers PlaceF (5’-GGGTAATCTTCTGAAAACGCATCGT-3’) from Hanif *et al*., 2013 and a reverse primer (5’-CTGGTCTCAGCGTCTGTTCA-3’) from the 5.8S region of the *H. perezi* rRNA gene cluster. The PCR was conducted on a BioRad C1000 thermocycler with AccuStart II PCR ToughMix under the following conditions: initial denaturation at 94°C for 5 minutes, followed by 31 cycles of 94°C for 25 seconds, 58°C for 25 seconds, and 72°C for 1 minute. Amplified products were visualized by 2% (w/v) agarose gel electrophoresis, stained with SYBR Safe DNA Gel Stain, and examined under ultraviolet light. Aliquots of each reaction were sequenced using an ABI 3130 genetic analyzer. Quality trimming and removal of primer sequences and low quality sequence was carried out using CLC Genomics Workbench 9.5.2 (Qiagen, Hilden, Germany). Assembled forward plus reverse assemblies were manually inspected, and consensus sequences were exported for further analysis.

Comparison of sequences derived from crabs in this study was conducted by alignment tools in CLC Workbench. A BLAST search of the 650 nt region identified candidate *H. perezi* sequence accessions in GenBank for comparison to experimental data. Selected candidates were chosen that represented genotypes I, II, and III (Small *et al*., 2012) and geographic origins of Europe, Asia and North America. The reulsting alignment was trimed to the sequence area in common, visually inspected, and visualized using the tree functions using https://ngphylogeny.fr/. Default parameters were used to create a Neighbor Joining tree with bootstrap values determined using 100 iterations.

#### Histological analysis

At the time of dissection for DNA and RNA extractions, pieces of hepatopancreas were excised from the crab samples and fixed for 24 hours in modified Davidson’s fixative, prepared with filtered seawater to maintain isotonicity. After fixation, the samples were transferred to 70% ethanol and stored at 4°C. Selected samples of fixed tissue were dehydrated in graded ethanol series, cleared in xylene and paraffin-embedded. Tissue sections were cut at 3-5 µm thickness, stained with Mayer’s hematoxylin and eosin (MHE) and coverslipped in NeoMount. Stained sections were inspected under a Leica DM3000 LED microscope. Images were captured with a Leica DMC 4500 camera and assembled and annotated in PhotoShop CS5 (Adobe Systems).

## Results

There was no evidence found for CsRV1 virus RNA in the samples of *C. sapidus* collected in this study, from the Aegean Sea, Adriatic Sea or Guadiana estuary. This represented a total of 166 individual samples from 6 locations. CsRV1 RT-qPCR assays were conducted on all RNA samples, but of these, several RNA samples (10 from Portugal and 4 from Greece), did not produce RT-PCR amplicons with the positive control SSU primers, and so were excluded from analysis.

DNA of the parasite *H. perezi* was detected in crabs from the Adriatic Sea and Guadiana estuary, but not from the Aegean Sea. The proportion of infected crabs and the mean intensity (copies of *H. perezi* target DNA mg^−1^ muscle tissue) are listed in Table 2. Individual crabs had *H. perezi* DNA intensity ranging from 30 copies to over 100,000 copies of target DNA mg^−1^ of tissue (supplemental material available).

**Table 2.** Prevalence and mean copy number of *H. perezi* and CsRV1 as determined by qPCR. Copy number of cloned target genes per mg tissue.

| Location | *Hematodinium perezii* |  |  | CsRV1 |  |
| --- | --- | --- | --- | --- | --- |
|  | Sample | Prevalence | Mean | Sample | Prevalence |

|  | size | (%) | intensity<br>(copies mg<sup>-1</sup>) | size* | (%) |
| --- | --- | --- | --- | --- | --- |
| Parila lagoon,<br>South Adriatic Sea | 30 | 23 | 1.28 x 10<sup>3</sup> | 30 | 0 |
| Ston saltworks, South<br>Adriatic Sea | 30 | 40 | 1.54 x 10<sup>5</sup> | 30 | 0 |
| Mlini,<br>South Adriatic Sea | 30 | 67 | 3.00 x 10<sup>5</sup> | 30 | 0 |
| Ada Bojana,<br>South Adriatic Sea | 30 | 10 | 3.72 x 10<sup>3</sup> | 30 | 0 |
| North Euboean Gulf,<br>Aegean Sea | 30 | 0 | n.a. | 26 | 0 |
| Foz de Odeleite,<br>Guadiana estuary | 30 | 87 | 3.25 x 10<sup>4</sup> | 20 | 0 |

\*some samples were excluded because they failed the rRNA amplification quality check.

Histological analysis of the hepatopancreas of a crab from Ada Bojana Montenegro that had 1.8 x10e3 gene copies mg^−1^ of fresh weight is shown in **Figure 2**. The interstitial spaces between tubules is distended and heavily invaded by parasite cells. Occasionally, cellular aggregates, possibly fixed phagocytes, are seen. Some of the tubules (T) are depleted of lipid inclusions (arrows).

**Figure 2.**
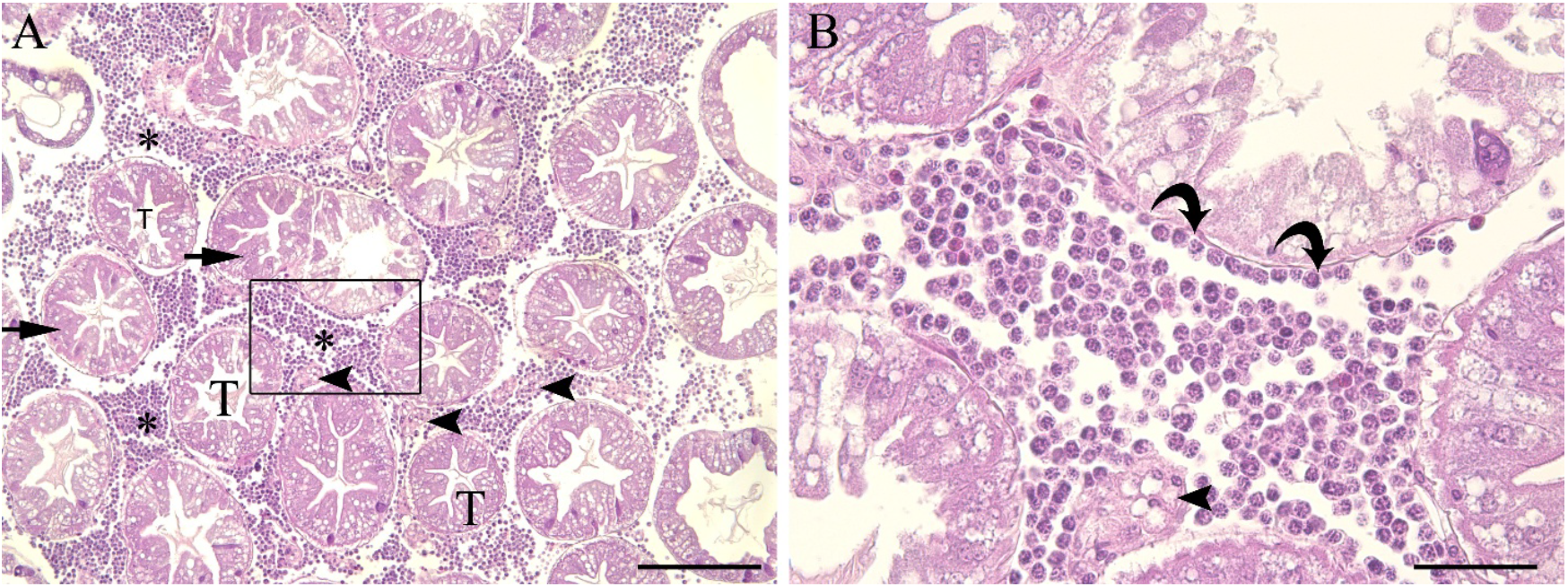
Representative micrographs of a hepatopancreas of the *C. sapidus* from Ada Bojana Montenegro infected with *H. perezi* at high infection intensity. **(A)** The intertubular space is distended and the interstitial tissue is almost completely replaced by vegetative stages of the parasite (asterisks). Occasionally, cellular aggregates, presumably fixed phagocytes, can be seen (arrowheads). Some of the tubules (T) show a depletion of lipid inclusions (arrows). **(B)** High magnification (400x) showing details of the developmental stages of the parasite with fixed phagocytes (arrowhead) and parasites adhering to the tubule epithelium (curved arrows). H&E staining. Scale bar: (A) 200 µm, (B) 50 µm

*H. perezi* infections (qPCR signal) in three locations were intense enough to enable amplification of a 888 nt region of the ITS and gene. Each of these sequences is deposited in GenBank, under the accession numbers PP854984 (Portugal), PP854699 (Montenegro), PP854700 (Zupa). As can be seen in **Figure 3**, a Maximum Likelihood phylogenetic analysis of aligned ITS1 regions of these sequences with corresponding regions of 256 nucleotides of the *H. perezi* ITS1 from other regions of the eastern Atlantic, North America, and Asia shows that the two *H. perezi* sequences from the Adriatic are identical, and slightly less similar to the sequence from Portugal (98% identity). All of them are more similar to *H. perezi* sequences from European waters than from Asia or North America. Bootstrap values are below 75 for all nodes except those at the branchpoints between continental regions.

**Figure 3.**
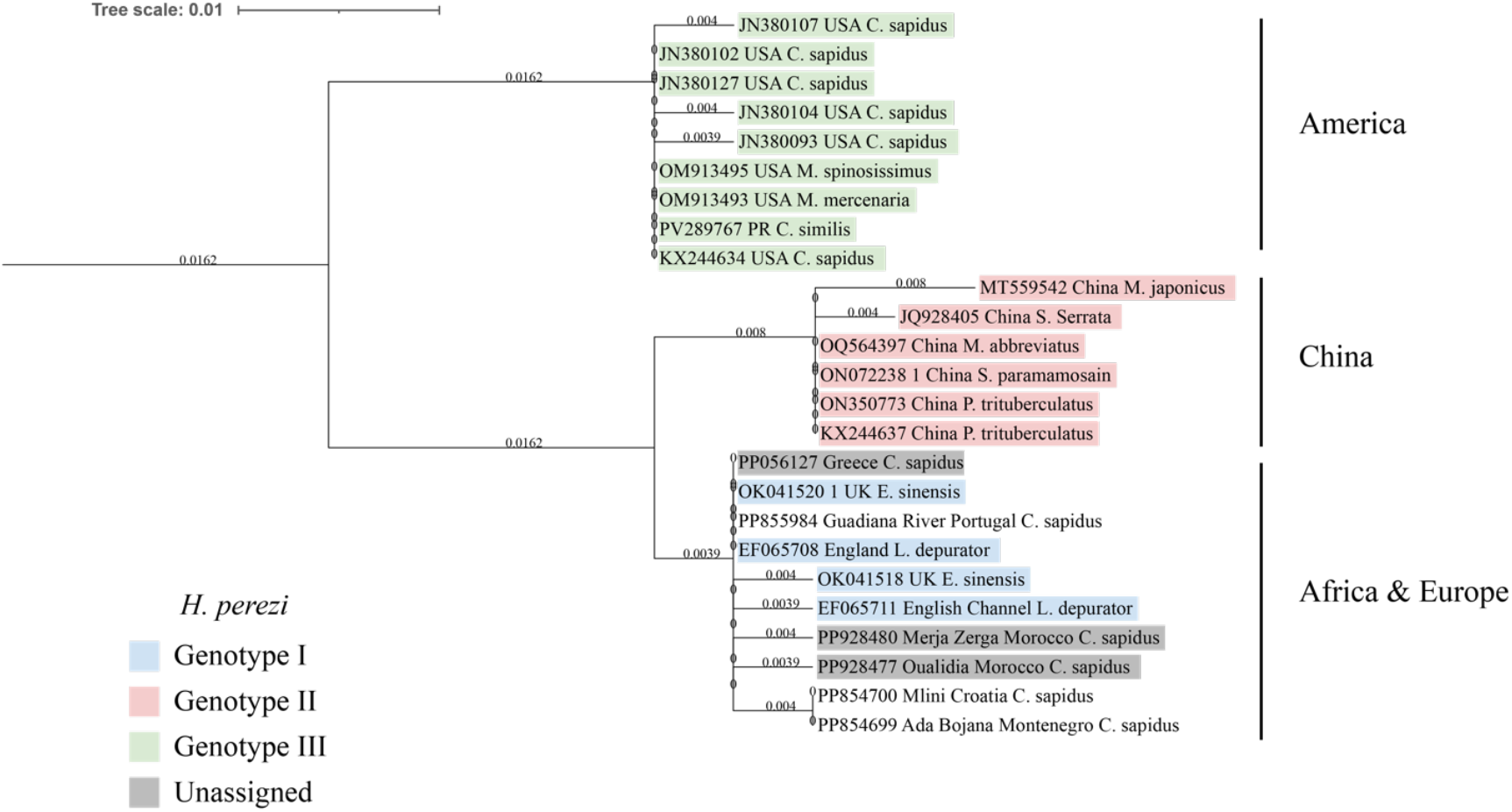
Neighbor joining tree from alignment (261 nt) of *H. perezi* ITS1 sequences from this study (no shading) as compared to *H. perezi* sequences in hosts from the Atlantic coast of Europe and North Africa, Mediterranean, Asia and North America. Numbers at branches are bootstrap values for 100 replicates. Genbank accession of *H. perezi* representing genotypes I, II and III are shaded with different colors as indicated in the legend.

## Discussion

The spread of *C. sapidus* throughout the Mediterranean and eastern Atlantic is highly consequential for the ecology and economy of the region. Measuring, predicting and adapting to the specific impacts of this invasive species is important for practical reasons, and presents a rare opportunity to evaluate ecological and evolutionary theories about invasive species escaping predators and disease. There has been some indication that blue crabs can dramatically decrease the population of aquatic prey (Céspedes *et al*., 2024; Nardelli *et al*., 2024). In Italy and Spain, bivalve farmers have abandoned the practice because of damage caused by *C. sapidus* predation (Chiesa *et al*., 2025). Often, invasive species exhibit a surge in abundance, followed by a decline and leveling of abundance in a new region after some years (Simberloff and Gibbons, 2004). In some cases this could be a consequence of the introduced species disrupting its own prey or habitat resources. Additionally, within the “enemy escape” framework/concept, this could be a result of both external predators and internal pathogens adapting to and exploiting the presence of the new species.

The current study gives insight to the presence of two known infectious agents of the Atlantic blue crab *C. sapidus* in selected locations of western Mediterranean and Atlantic Iberian estuaries. The opportunity to analyze crabs from the Aegean Sea, Adriatic, and Guadiana estuary represents an east-to-west range that corresponds to the possible expansion trajectory of the species over time (Mancinelli *et al*., 2021). The two infectious agents studied represent very different life history strategies for pathogens. The CsRV1 virus is described only in the *C. sapidus* native range, and has not been yet been reported in any other species of host (Zhao *et al*., 2022). In contrast, the protozoan parasite (parasitic dinoflagellate) *H. perezi* is a globally-distributed crustacean host generalist (Small and Li 2024).

There are relatively few studies related to the health status of *C. sapidus* in the Mediterranean and adjacent waters. At the time of sampling, ours was the first study to investigate the virus CsRV1 in the Mediterranean or eastern Atlantic. Although we did not detect CsRV1 in any of the *C. sapidus* tested, our limited study cannot be taken as evidence that the virus is entirely absent from the Mediterranean or eastern Atlantic. Perhaps the most intriguing suggestion that CsRV1 could be present in the Mediterranean is the report by Walton *et al*., (1999) on the sequence of a segment of the P virus genome in the portunid crab *Liocarcinus depurator*. The segment of P virus has over 95% identity to segment 4 of CsRV1 (Flowers *et al*., 2016). This level of identity is in the range of genome difference between CsRV1 ecotypes in north America versus south America (Zhao *et al*., 2020). Future investigations of CsRV1 in the Mediterranean should include assessments of portunid species such as *Liocarcinus* sp. and *Carcinus* sp.

On the North American Atlantic coast, studies have shown that CsRV1 prevalence is highly variable across time and space, often as low as 0% or as high as 75% prevalence (Flowers *et al*., 2018). The only consistent trend seen for CsRV1 in the native range is the near absence in the tropics (Zhao *et al*., 2020). Flowers *et al*., (2018) noted that high prevalence of CsRV1was coincident with higher *C. sapidus* populations in Chesapeake Bay, which is consistent with the principles of host-dependent prevalence (Cortez and Duffy 2021). All of the locations surveyed in the current study are temperate, but the overall blue crab population numbers are also unknown in each of the areas. In the coming 5 – 10 years, it will be informative to re-survey for the presence of CsRV1 virus in areas with particularly high *C. sapidus* populations or where the species has been recorded for a long time.

The presence of *H. perezi* in Mediterranean *C. sapidus* was first reported by Mancinelli *et al*., (2013) in ecosystems of the Apulia coastline. In that study, over 70% of the crabs examined (*C. sapidus, Eriphia verrucosa, Pachygrapsus marmoratus* and *Carcinus mediterraneus*) in the Torre Colimena (Ionean Sea) basin and Acquatina lagoon (West Adriatic Sea) were infected by *Hematodinium* sp. (Mancinelli *et al*., 2013). Our study showed that just across the southern Adriatic, prevalence of *H. perezi* was moderate to high at all locations. These two coasts are highly connected by an anticlockwise gyre in the southern (Estournel *et al*., 2021), which would have the potential to rapidly convey both crab larvae and parasite propagules between them.

In the Aegean Sea, Lattos *et al*., (2024) recently reported 85% prevalence of *H. perezi* in *C. sapidus* that were collected in November following 2020 mortality events. They suggest this high prevalence is responsible for recent declines in *C. sapidus* abundance in that region of Greece.

These authors pointed out that there were no external signs of disease observed during the sampling of *C. sapidus* but that infected crabs had a characteristic “milky” coloration of hemolymph. In our study of Aegean Sea crabs harvested in April of 2021, we detected no *H. perezi*. Our collection was from a local fish market, so lacked fine scale specificity of geographic origin. We are aware that the use of commercially harvested crabs may bias against heavily infected or moribund crabs, the work of Lattos *et al*. (2024) reported that both “moribund and healthy individuals” were found with *H. perezi*. The association of crab declines with *H. perezi* is still relevant to our current study; there are anecdotal reports from fishermen in Mlini, which had the 2^nd^ highest prevalence (76%), that they didn’t encounter blue crabs after the summer of 2021 for almost a year (M. Pećarević, personal communication). While there are many possible reasons for this, one is that infections by *H. perezi* reduced the abundance of blue crabs, similar to what has been reported in the *C. sapidus* native range (Lee & Frischer 2004).

On the Atlantic coast of north Morroco, Lamkhalkhal *et al*., (2024) also recently reported *H. perezi* in *C. sapidus* at 25-60% prevalence. Moreover, they show that the sequence of the ITS1 gene of *H. perezi* from Morocco is similar to that in strains from Europe and the Mediterranean. Our study is the first in our knowledge to document *H. perezi* on the Atlantic coast of Portugal. We are aware that a 2021 metagenomic investigation of *C. sapidus* stomach contents from the Rio Formosa estuary near Faro, Portugal also revealed evidence of *H. perezi* DNA. In this case, the sequence was a match to the mitochondrial COX1 gene (J. Encarnacao and P. Morais, personal communication), so no direct comparison can be made to the ITS1 sequences. Nevertheless, this suggests that the parasite has been in Atlantic coastal estuaries of Portugal since at least 2021.

In the *C. sapidus* native range, the prevalence of *H. perezi* is typically associated with high salinity (Messick and Shields 2000; Gandy *et al*., 2015). In our study, there is no strict correspondence between salinity at the sites sampled and the prevalence of *Hematodinium* sp. In fact, the highest prevalence was found in the Guadiana River, which had locally low salinity but is less than 20 km away from the ocean. In the Eastern Adriatic Sea, salinity was above 30 psu except in Parila lagoon (18 psu), which also had the lowest *Hematodinium* sp. prevalence in that region. The only sample without detection of *H. perezi* was in the Aegean Sea, where crabs were obtained from a fish market. Therefore precise location and salinity information was not available. This is in contrast to the recent report by Lattos *et al*., (2024) of *H. perezi* prevalence over 80% across the Aegean Sea, which they note was coincident with a decline in the abundance of *C. sapidus* as reported by fishermen. The contrasting prevalence found in our study and that of Lattos *et al*., (2024) is not unusual for marine disease studies, where diverse environmental factors interact with shifting population size and connectivity to produce highly variable prevalence.

The relationship of *H. perezi* ITS1 sequences in our study with corresponding regions of *H. perezi* strains across the globe is consistent with prior observations that *H. perezi* within different continents are similar to one another, and distinct from other regions, regardless of host. The high identity of ITS1 sequences between Greece and Portugal strains stands out, however, and may suggest a rapid movement from one location to another. One possible mechanism for such transport of a parasite strain is ballast water (Nehring, 2011; Cantrell *et al*., 2020). To comply with the International Convention for the Control and Management of Ships’ Ballast Water and Sediments (IMO, 2004), prior to entering the Mediterranean, cargo ships exchange ballast water in the Atlantic. Therefore, it is possible that a strain of *H. perezi* or a host infected with it was transferred in ballast from the Atlantic to the Aegean Sea. Additional comparisons of *H. perezi* genetics will be needed to lend support or refute the idea that that the parasite has recently moved.

It is important to understand how endemic infectious agents find or adapt to newly arrived invasive species for at least two reasons. One is the effect that these agents may have on the health of native species; will they proliferate in a new reservoir host, or perhaps the new host will constitute a dead end that lowers transmission of the infectious agent. Evidence is that the genotype I of *H. perezi* is well-adapted to *C. sapidus*. A second reason to understand infectious agents is the impact on the invasive species over time. In many regions of Europe, there is still a policy to reduce the number of *C. sapidus* in the invaded range. As income from *C. sapidus* fisheries grow, however, there may be concern about disease-related mortality if it affects harvests. In North America, the CsRV1 virus is associated with stress and high populations of *C. sapidus* (Spitznagel *et al*., 2019; Flowers *et al*., 2018). Elsewhere in the Americas, other viruses have been recently identified in *C. sapidus* (Tavares *et al*., 2022; Zhao *et al*., 2022), illustrating that there is a potential for various pathogens to be transferred to or within European waters by ballast water transfer or movement of live crustaceans. Surveillance of *C. sapidus* health is therefore advisable in all regions of Europe.

## Funding and Acknowledgements

OP received support from the Smithsonian Environmental Research Center. ES and OP were partially funded by the National Oceanic and Atmospheric Administration, Office of Education Educational Partnership Program award number (NA21SEC4810005). Its contents are solely the responsibility of the award recipient and do not necessarily represent the official views of the U.S. Department of Commerce, National Oceanic and Atmospheric Administration.

## Statements and Declarations

The authors have no financial or non-financial interests that are directly or indirectly related to the work submitted for publication.

